# MicroRNA miR-100 decreases glioblastoma growth by targeting SMARCA5 and ErbB3 in tumor-initiating cells

**DOI:** 10.1101/865105

**Authors:** Bahauddeen M. Alrfaei, Raghu Vemuganti, John S. Kuo

## Abstract

Glioblastoma multiforme (GBM) is the most aggressive and most frequently diagnosed malignant human glioma. Despite the best available standard of care (surgery, radiation, and chemotherapy), the median survival of GBM patients is less than 2 years. Many recent studies have indicated that microRNAs (miRNAs) are important for promoting or reducing/limiting GBM growth. In particular, we previously showed that GBMs express decreased levels of miR-100 relative to control tissue and that restoring miR-100 expression reduced GBM tumorigenicity by modulating SMRT/NCOR2 (Nuclear Receptor Corepressor 2). Here, we demonstrate that miR-100 overexpression decreases expression of the stem cell markers, nestin and L1CAM, and decreases proliferation of GBM tumor-initiating cells (GBM stem-like cells). We further show that miR-100-mediated anti-tumorigenic activity limits the activity of SMARCA5 and its downstream target STAT3 (known as mTOR-STAT3-Notch pathway). In addition, we report ErbB3 (Her3) as a putative miR-100 target, including inhibition of its downstream AKT and ERK signaling pathways.

## Introduction

Glioblastoma multiforme (GBM), the most aggressive primary brain tumor, accounts for more than 50% of all detected malignant brain cancers and approximately 20% of all primary intracranial tumors[1, 2]. Approximately 15,000 new cases of GBM and CNS malignancies are diagnosed annually in the USA [3]. Median patient survival is under 2 years even with the best standard of care [4, 5]. The molecular mechanisms responsible for GBM growth and invasion are poorly understood.

MicroRNAs (miRNAs) are small, non-coding RNAs (16–22 nucleotides long) that repress protein translation by binding to the 3’UTRs of mRNAs [6, 7]. Many miRNAs are known to be differentially expressed in various types of cancer [8]; it is therefore essential to understand their role in tumorigenesis [9]. At least 19 miRNAs have been identified as being linked to the pathogenesis of GBM [10, 11]. Of these, miR-100 has been linked to several targets that are known to modulate GBM growth and/or survival, such as fibroblast growth factor receptor 3 (FGFR3), silencing mediator of retinoic acid and thyroid hormone receptor (SMRT), and ATM Serine/Threonine Kinase (ATM; ataxia telangiectasia mutated) [12-14]. Moreover, miR-100 was reported to stimulate a positive therapeutic response in breast cancer stem-like cells undergoing hormonal therapy [15].

We previously demonstrated that downregulation of miR-100 promotes GBM growth and invasion, and that restoring miR-100 expression reduces GBM growth and survival [13]. We also showed that miR-100 limits GBM tumor proliferation and extends the survival of mice bearing orthotopic GBM xenografts by inhibiting the miR-100 target, SMRT/NCOR2 [13]. These findings were subsequently supported by Luan *et al*. (2015), who also confirmed the tumor suppressor activity of miR-100 and examined two GBM cell lines (U251 and T98G) in addition to 13 patient GBM specimens. They confirmed the decrease in endogenous levels of miR-100 in all tested tumor samples and in both cell lines, and compared GBM tumors to adjacent normal tissue. When miR-100 was overexpressed in GBM cell lines, reduced proliferation, migration, and chemo-sensitivity were observed. The authors concluded that miR-100 has anti-tumor activity against GBMs.

Since a single miRNA can regulate multiple target mRNAs, the beneficial effects of miR-100 might also be mediated by other target proteins in addition to SMRT/NCRO2. In this study, we have uncovered and evaluated additional pathways that might also be regulated by miR-100 and potentially contribute to its anti-tumorigenic activity. In particular, we evaluated the role of ErbB family members that we found are targeted by miR-100. Members of this protein family are known to be resistant to EGFR inhibitor therapy especially in GBM stem-like cells [16]. Two members of the ErbB family Her2 (ErbB2) and Her3 (ErbB3) are being targeted by immunotherapy in clinical trials designed to control GBM growth [17, 18].

In this study, we restored miR-100 levels in primary GBM cells by transfecting GBM tumor-initiating cells (TICs) with pre-miR-100, which generates two forms of miR-100: miR-100-5p and miR-100-3p. We found that the dominant isoform, miR-100-5p [19], is down-regulated in GBM-derived TICs (also called GBM stem-like cells) known to be responsible for tumor progression and recurrence [19] because of radiation and chemo-resistance [20]. Thus, in this study, we tested the therapeutic utility of miR-100-5p overexpression for controlling GBM growth and also evaluated the downstream mechanism of miR-100-5p activity.

## Materials and Methods

### Isolation and validation of GBM TICs

All human tumor specimens were collected after patient informed consent and with approval of University of Wisconsin-Madison Institutional Review Board. Patient-derived TICs were isolated from GBMs and validated as previously described [16, 21-23]. Tumor tissue collected from the operating room was minced and chopped twice using a tissue chopper (Sorvall TC-2 Smith-Farquhar) and plated in medium (70% Dulbecco’s modified Eagle medium-high glucose, 30% Ham’s F12, 1X B27 supplement, 5 μg/ml heparin, penicillin-streptomycin-amphotericin, and 20 ng/ml each of epidermal growth factor (EGF) and basic fibroblast growth factor (bFGF)). We used three primary (22, 33, and 44) GBM samples and one recurrent GBM sample (12.1). All of the GBM TICs were negative for EGFRvIII mutations [16]. A tumor-free neural stem cell (NSC) line prepared from fetal human cortical tissue and used as a source of normal control cells was a kind gift of Dr. Clive Svendsen [23].

### Real-time (Quantitative) PCR

Total RNA was isolated using an RNA isolation kit (Life Technologies). All probes and primers were purchased from Life Technologies, USA, and quantitative PCR was conducted using TaqMan assays according to manufacturer’s instructions (Life Technologies, USA). 18s rRNA was used as a house-keeping control as described previously [16].

### miRNA and siRNA transfection

Cells were transfected with miRNA precursors, siRNAs, miRNA mimics and the miR-100 isoforms miR-100-5p and miR-100-3p (all from Life Technologies) using 15 pmol PepMute reagent (SignaGen Labs, Rockville MD) per million cells as described previously ^19,20^.

### Cell viability assay

GBM TICs were grown as spheres, disseminated into single cells, then inoculated in 96-well plates at a density of 20,000 cells/well and temperature of 37°C. After 1 day of growth, the cells were transfected with pre-miR-100 or with a control miR. Following 2 days of growth in culture medium containing 4.5g glucose DMEM/F12, 20 pg EGF and 20 pg bFGF, cell numbers were quantified using an MTS assay (CellTiter 96 Aqueous, Promega, USA) as per the manufacturer’s instructions.

### Cell proliferation assay

The Click-iT EdU assay (which is similar to the BrdU assay) was performed according to the manufacturer’s (Invitrogen) instructions. Twenty thousand cells were plated and then transfected with pre-miR-100 or control miRs in combination with the SMRT/NCOR2 expression vector (pSMRT, Fisher Scientific) after 1 day of growth. The Click-iT EdU assay was performed following an additional 2 days of growth in culture medium containing DMEM/F12, EGF and bFGF.

### Luciferase reporter assay

293T cells were co-transfected with pre-miR (15 pmol) and a luciferase reporter plasmid (1 ug) containing the 3’UTR of SMARCA5 or ErbB3 mRNA in 96-well culture plates. Twenty-four to forty-eight hours after transfection, the light switch luciferase assay (Switch Gear Genomics, Menlo Park, CA) was performed according to the manufacturer’s instructions. The signal was detected using a microplate luminometer (Turner Biosystems, Inc., CA) running Veritas software version 1.9.2. The reporter plasmids contained the 3’UTRs of SMARCA5 or ErbB3 mRNA, which include the miR-100 seed sequence. The control plasmids (negative controls) contained a mutated seed sequence.

### Western blotting

Western blotting was performed as previously described [16, 24]. Cell lysates were collected, and the protein concentration was determined by the Bradford assay (Bio-Rad, CA). The protein samples were electrophoresed on SDS-PAGE gels, transferred to PVDF membranes, and analyzed using antibodies against α-tubulin, phospho-AKT (S473), total-AKT, phospho-ERK (Thr202/Tyr204), total-ERK, phospho-STAT3 (Tyr705), STAT3, p21, SMARCA5 (Santa Cruz Biotechnology, USA), SMRT/NCOR2 (Santa Cruz Biotechnology, USA), ErbB3 (Santa Cruz Biotechnology, USA), ErbB2 (Santa Cruz Biotechnology, USA), human nestin (Santa Cruz Biotechnology, USA), and L1CAM (Fisher, USA). All antibodies were purchased from Cell Signaling, USA, unless otherwise indicated.

### Tumor xenograft assay

UW-Madison institution-approved animal protocol was followed for all experimental procedures. Tumor xenografts were generated via stereotactic implantation of tumor cells as described previously [13]. Briefly, GBM cells were enzymatically dissociated into single cells. One million cells were suspended in 5 μl of PBS and stereotactically implanted at 0.33 μl/min into the right striatum of anesthetized immunodeficient 6-8 weeks old NOD-SCID mice (Jackson lab) at the following coordinates referenced from bregma: 0 mm anteroposterior, +2.5 mm mediolateral, and −3.5 mm dorsoventral [25]. Xenograft growth was detected and verified by MRI, and brains containing the xenografts were obtained from the animals after death.

### Inducible and stable expression of pre-miR-100 and immunohistochemical analysis

The 22T and U87 GBM cell lines were orthotopically implanted into immunodeficient NOD-SCID mice. Control and miR-100 doxycycline inducible overexpression vectors (GeneCopoeia, MD) were created and then integrated into the genomes of tumor cells with lentiviruses. Later, at least one million cells were implanted into mouse brains. The vectors resulted in a two- to three-fold increase in expression above baseline miR-100 levels. Those vectors were validated and carried out as described as previously published [13]. When severe clinical symptoms were detected, or the mice were moribund, they were sacrificed. The brains of the mice were formalin-fixed, paraffin-embedded, sectioned (into 5 µm-thick sections), and stained with hematoxylin, as described previously [26].

### Statistics

The statistical analyses were performed using Student’s t-test and one-way ANOVA/Tukey’s multiple comparison post-tests. All error bars represent the standard error of the mean (S.E.M.), and the significance level (*) was P < 0.05.

### Institutional approval

All of the procedures involving human specimens were approved by the University of Wisconsin-Madison Human Subjects Institutional Review Board, and informed consent was obtained from patients. In addition, all animal procedures were approved by the UW-Madison Institutional Animal Care and Use Committee.

## Results

### Expression of miR-100-5p is down-regulated in GBM cells

Quantitative PCR revealed that expression of endogenous miR-100-5p was reduced by approximately 50–80% in the four GBM tumor cell populations (12.1, 22, 33, and 44) evaluated relative to the levels of endogenous miR-100-5p in two different control normal human neural stem cell lines (n = 3/group; P<0.05; Fig. 1A).

**Figure 1:**
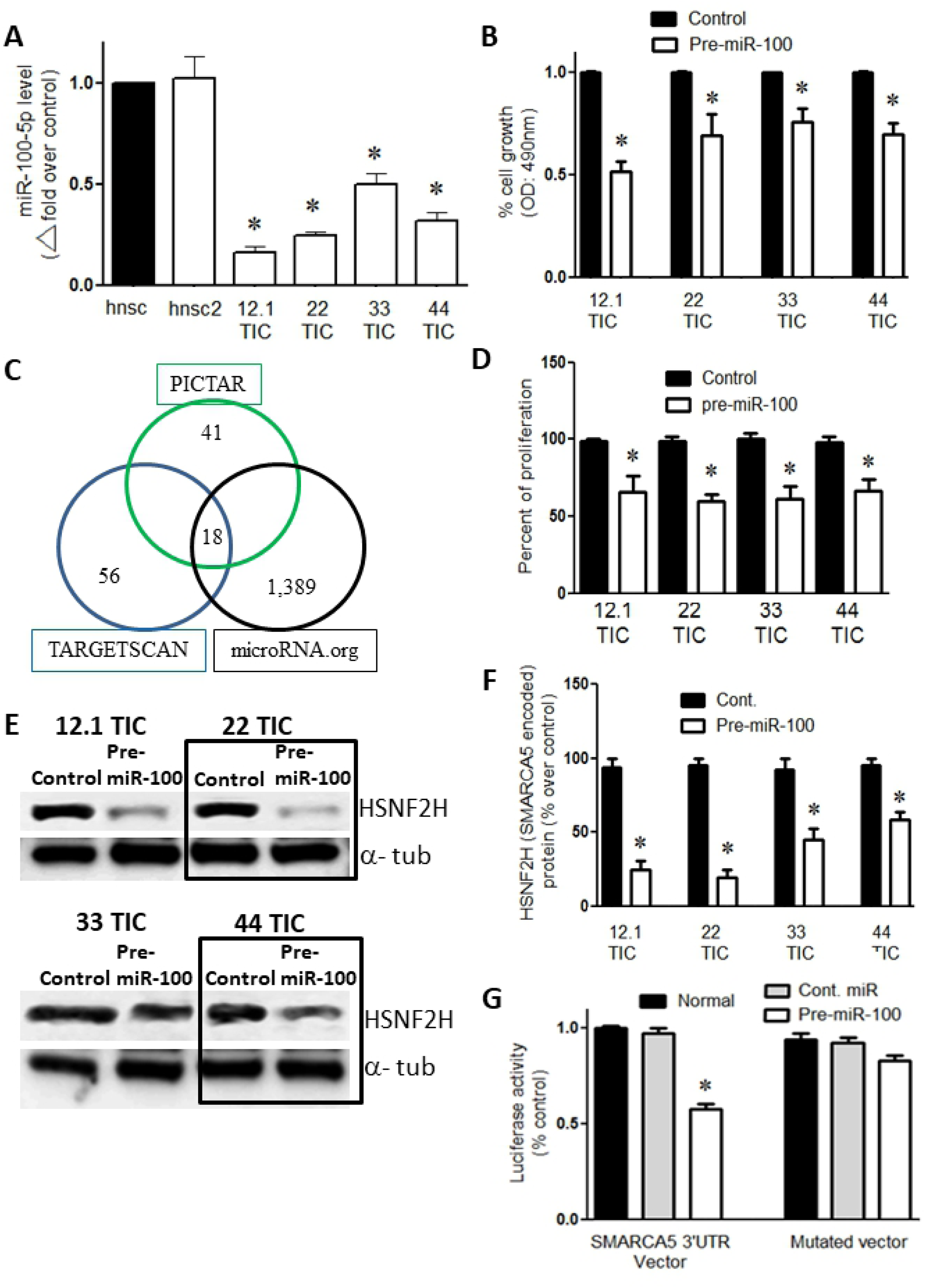
Overexpression of pre-miR-100 reduces proliferation and targets SMARCA5. (A) qPCR shows that the expression of miR100-5p is lower in GBM TICs (12.1, 22, 33, and 44) compared to human neural stem cells lines (hnsc and hnsc2). (B) The overexpression of pre-miR-100 reduces the number of GBM TICs compared to controls as determined by the MTS assay. (C) The targets of miR-100-5p as predicted by three algorithms: PICTAR, TARGET SCAN, and microRNA.org. The intersection of the circles represents the number of shared targets. (D) The overexpression of pre-miR-100 decreases proliferation in GBM TICs compared to the control. (E) Immunoblots showing the reduction of SMARCA5-encoded HSNF2H protein levels following the transient expression of pre-miR-100. The reduction in HSNF2H levels does not occur when control miR is overexpressed. The boxes are included to make the blot easier to interpret. (F) The quantification of the immunoblot in panel (E) shows that HSNF2H levels are reduced by 40–70% when pre-miR-100 is overexpressed. (G) Pre-miR-100 inhibits the luciferase signal from the SMARCA5 3’UTR reporter compared to controls that utilize a different miRNA or a reporter with a mutated target site.

### Restoring miR-100 levels decreased cell viability and proliferation

When the TIC cell lines were transiently transfected with pre-miR-100, a 20–50% decrease in cell viability was observed (n = 3/group; p<0.05; Fig. 1B) and a 20–40% decrease in cell proliferation (n = 3/group; p<0.05; Fig. 1D) compared to the respective controls transfected with a control miRNA.

### SMARCA5 mRNA is a target of miR-100-5p

Using three target identification algorithms (PICTAR, TARGETSCAN, and microRNA.org), we found 18 common putative targets of miR-100-5p (Fig. 1C). *SMARCA5* was the only common target that was among the top six targets identified by all three algorithms (Table 1). When the TICs (12.1, 22, 33, and 44) were transfected with pre-miR-100, the protein levels of HSNF2H (encoded by *SMARCA5*) were approximately 40–70% lower than observed in the control transfected cell lines (p< 0.05; n = 3; Fig. 1E, F). We further confirmed the miR-target relationship by co-expressing a SMARCA5 3’UTR luciferase reporter vector with pre-miR-100 or with miR-100-5p. When 293T cells were co-transfected with the SMARCA5 3’UTR and with either pre-miR-100 or miR-100-5p, SMARCA5 3’UTR luciferase activity was inhibited by 45% (p<0.05; n = 3) compared to cells transfected with the control miRNA (Fig. 1G).

**Table 1.** Predicted targets of miR-100. A list of the top six predicted targets of microRNA-100-5p (miR-100-5p) and their transcript ID numbers as they rank according to three different algorithms: microRNA.org, PICTAR, and TARGETSCAN. The asterisk (*) denotes targets that have the same score and thus can be given the same rank as the highest ranked candidate. Shading represents repeated appearance of SMARCA5.

| Rank | MicroRNA.org |  | PICTAR |  | TARGETSCAN 5.2 |  |
| --- | --- | --- | --- | --- | --- | --- |
| 1 | TMPRSS13 (NM_001077263) |  | SMARCA5 (NM_003601) |  | THAP2 | (NM_031435) |
| 2 | SMARCA5 (NM_003601) |  | BAZ2A | (NM_013449) | KBTBD8 | (NM_032505) |
| 3 | ANKAR | (NM_144708) | HS3ST3B1 (NM_006041) |  | HS3ST3B1 | (NM_006041) |
| 4 | ICK* | (NM_014920) | HS3ST2 | (NM_006043) | HS3ST2 | (NM_006043) |
| 5 | AP1AR* | (NM_018569) | FRAP1 | (NM_004958) | CTDSPL | (NM_005808) |
| 6 | NCOR2* | (NM_001077261) | EIF2C2 | (NM_012154) | SMARCA5 (NM_003601) |  |

### miR-100-5p-mediated inhibition of cell proliferation involves activation of AKT, ERK, and ErbB3

In the cell lines treated with pre-miR-100, there was a significant reduction in the phosphorylation of AKT and ERK relative to the cell lines transfected with the control pre-miRNA (Fig. 2A-D; p<0.05; n = 3/group). Treatment with siRNAs that are specific to SMRT or SMARCA5 also significantly inhibited AKT and ERK phosphorylation (Fig. 2A-D; p<0.05; n = 3). Furthermore, the knockdown of ErbB (ErbB functions upstream of AKT and ERK) significantly suppressed the phosphorylation of AKT and ERK in TICs (Fig. 2A-D).

**Figure 2:**
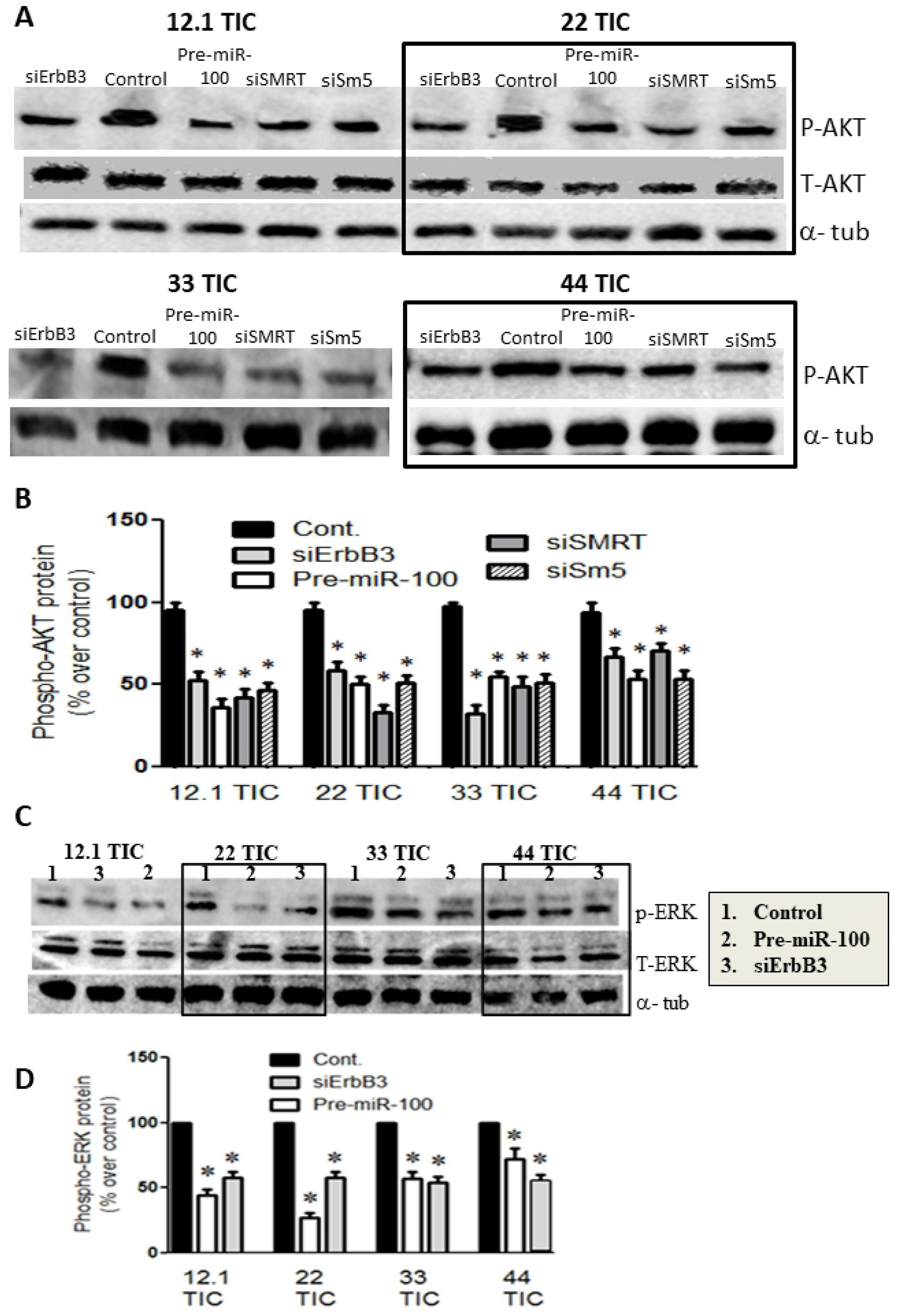
The activity of the AKT and ERK pathways is reduced following the overexpression of pre-miR-100. (A) Immunoblot showing the reduction in phospho-AKT protein levels following the transfection of GBM TICs with pre-miR-100 or with the following siRNAs: siErbB3, siSMRT, or siSm5 (siSMARCA5). This pattern is not observed in GBM TICs transfected with control miRNA. The boxes are included to make the blot easier to interpret. (B) Quantification of protein levels in panel (A) reveals that their reduction ranges from 20% to 60%. (C) Immunoblot showing the inhibition of phospho-ERK following the transfection of GBM TICs with pre-miR-100 or siErbB3. No inhibition of phospho-ERK is observed in the control. The boxes are included to make the blot easier to interpret. Total ERK (T-ERK) was measured to ensure that effect is only at phosphorylation level and not total protein. (D) Quantification of the protein levels in panel (C) shows that the reduction of phospho-ERK ranges from 10% to 70%.

Pre-miR-100 releases 2 mature miRNAs, miR-100-5p and miR-100-3p; our bioinformatics analysis showed that the miR-100-3p targets ErbB3 (Fig. 3C). We found that expression of miR-100-3p was 40–80% lower in GBM TICs than in control NSCs (Fig. 3D; p<0.05; n = 3). A luciferase reporter assay showed that miR-100-3p significantly inhibited the expression of the ErbB3 3’UTR vector (Fig. 3E; P<0.002; n = 3).

**Figure 3:**
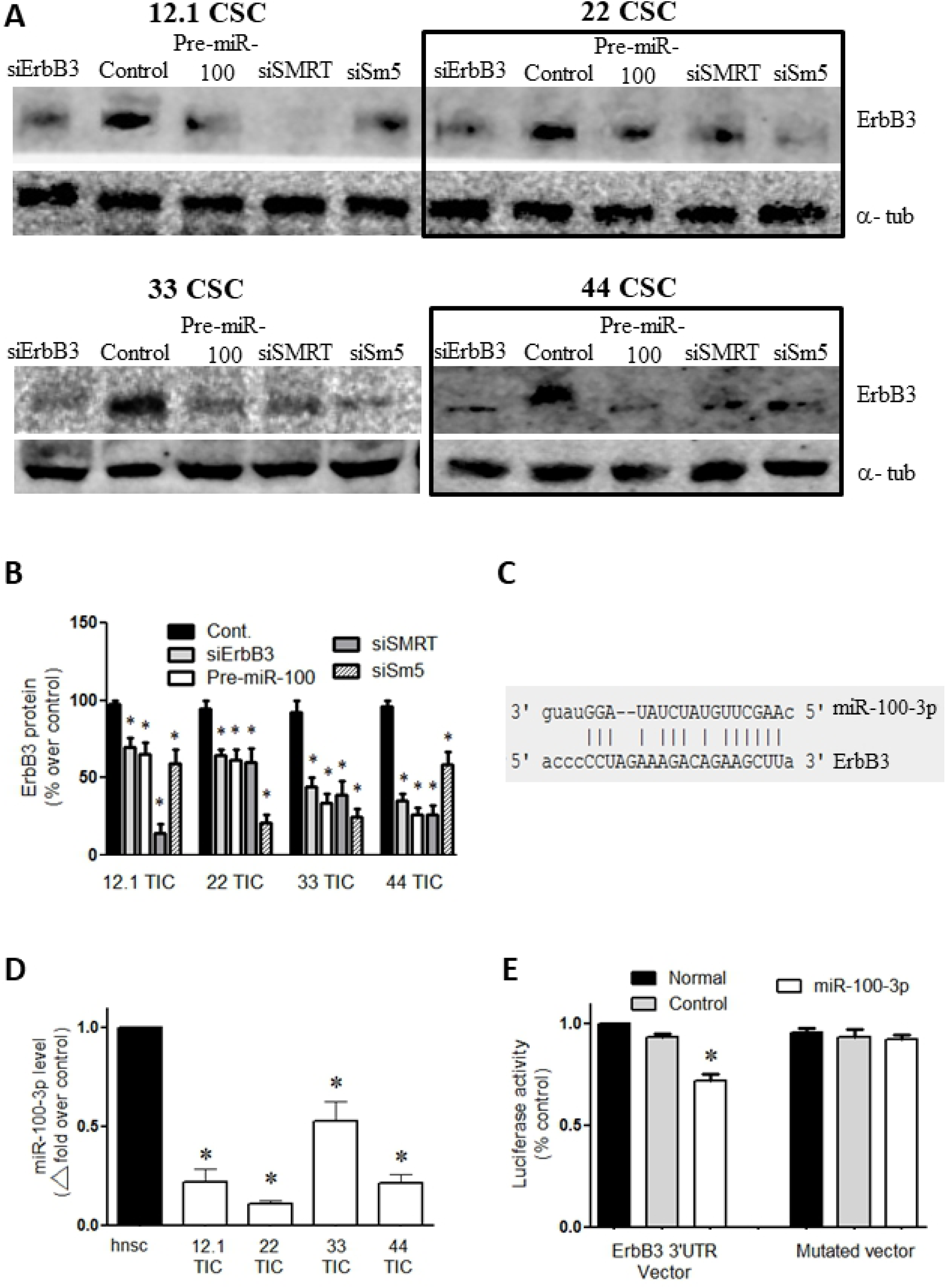
Overexpression of pre-miR-100 targets ErbB3 mRNA and decreases ErbB3 protein levels. (A) Verification of ErbB3 inhibition following transfection with pre-miR-100, siErbB3, siSMRT, and Sm5 (siSMARCA5). The boxes are included to make the blot easier to interpret. (B) Quantification of panel (A) shows a 30–80% reduction in ErbB3 (Her3) protein levels following transfection with pre-miR-100, siErbB3, siSMRT, or Sm5 (SMARCA5 siRNA). The control does not reduce ErbB3 expression. (C) Diagram showing the predicted binding of miR-100-3p to ErbB3 mRNA. (D) Internal expression of miR-100-3p in GBM TICs (12.1, 22, 33, and 44) compared to human neural stem cells (hnsc) as detected by qPCR. (E) miR-100-3p lessens the signal from the ErbB3 3’UTR luciferase reporter compared to controls that utilize a different miRNA or a reporter with a mutated target site. Asterisk denotes statistical significance of p<0.05.

### SMRT and SMARCA5 controls STAT3

When the GBM TIC lines were transfected with pre-miR-100, SMRT siRNA, or SMARCA5 siRNA, phosphorylation of STAT3 phosphorylation (a marker of STAT3 activation) was completely inhibited in all four of the tested cell lines (Fig. 4A, B; p<0.05; n = 3).

**Figure 4:**
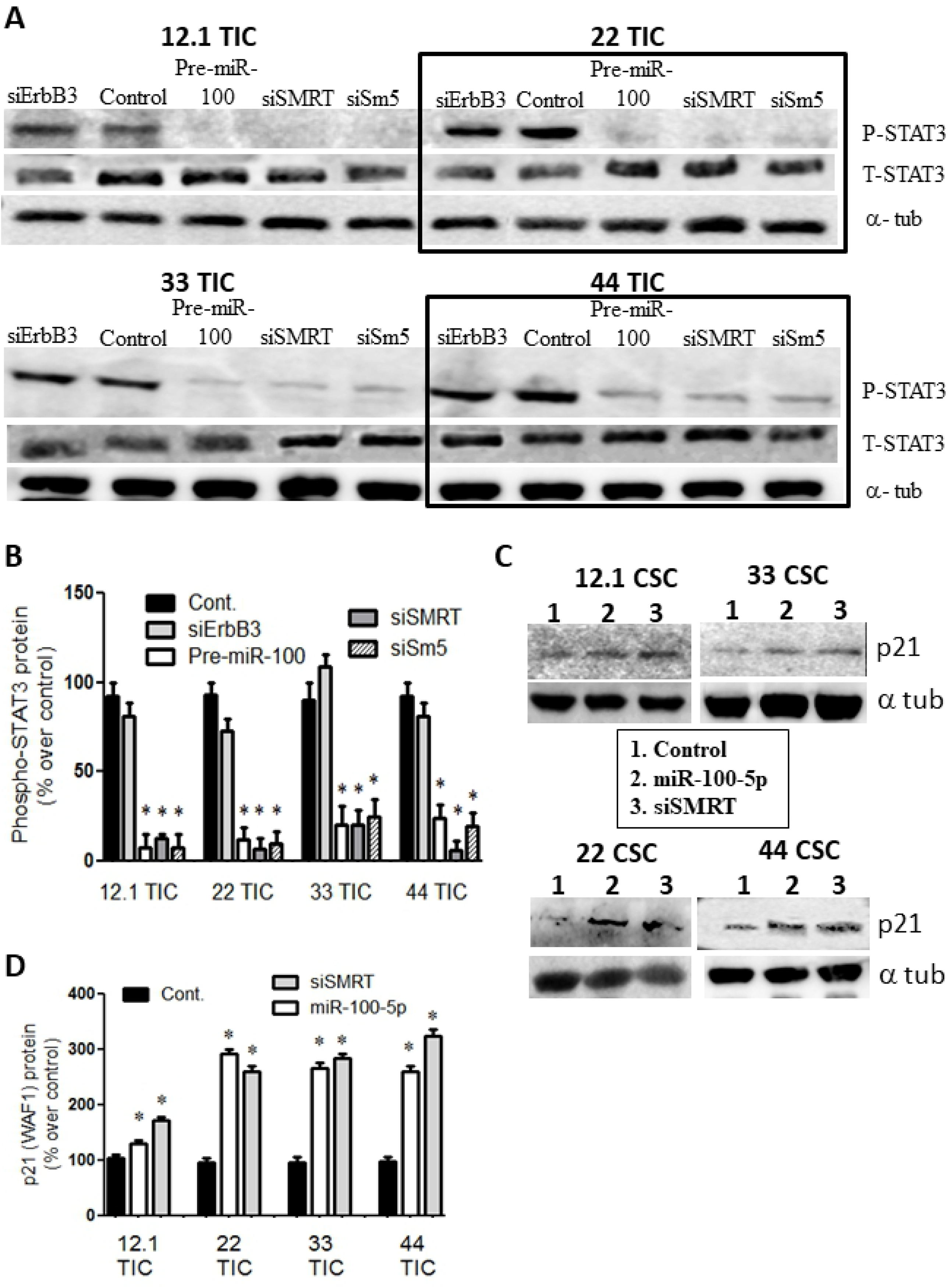
Overexpression of pre-miR-100 reduces phospho-STAT3 and increases p21 (WAF1). (A) Immunoblot showing a reduction in phospho-STAT3 when GBM TICs are transfected with pre-miR-100, siErbB3, siSMRT, or Sm5 (SMARCA5). This pattern is not observed following transfection with the controls. The boxes are included to make the blot easier to interpret. Total STAT3 (T-STAT3) shows no change in total protein level. (B) Quantification of the protein levels in panel (A) shows a reduction 60–90% of phospho-STAT3 except for the control and siErbB3. (C) p21 protein levels are elevated in response to miR-100-5p overexpression or SMRT silencing. (D) Quantification of the protein levels in panel (C) shows elevated p21 levels compared to the control by 20–300%. Asterisk denotes statistical significance of p<0.05.

### Overexpression of miR-100-5p upregulates p21

Western blot analysis showed that expression of the cell cycle inhibitor p21 (WAF1 or cyclin-dependent kinase inhibitor 1) was upregulated by 50–300% when the various GBM TICs were treated with pre-miR-100, miR-100-5p, or SMRT siRNA (Fig. 4C, D; P<0.05; n = 3).

### pre-miR-100 treatment decreases stem cell markers

Nestin and L1CAM are both stem cell markers that are normally expressed by GBM TICs. When the GBM TICs were transiently transfected with pre-miR-100 or with miR-100-5p, there was a 40–75% decrease in nestin levels (Fig. 5A, B; P<0.05; n = 3) and a 60% decrease in L1CAM levels (Fig. 5C, D; p<0.05; n = 3) compared to the control miRNA-transfected cell lines.

**Figure 5:**
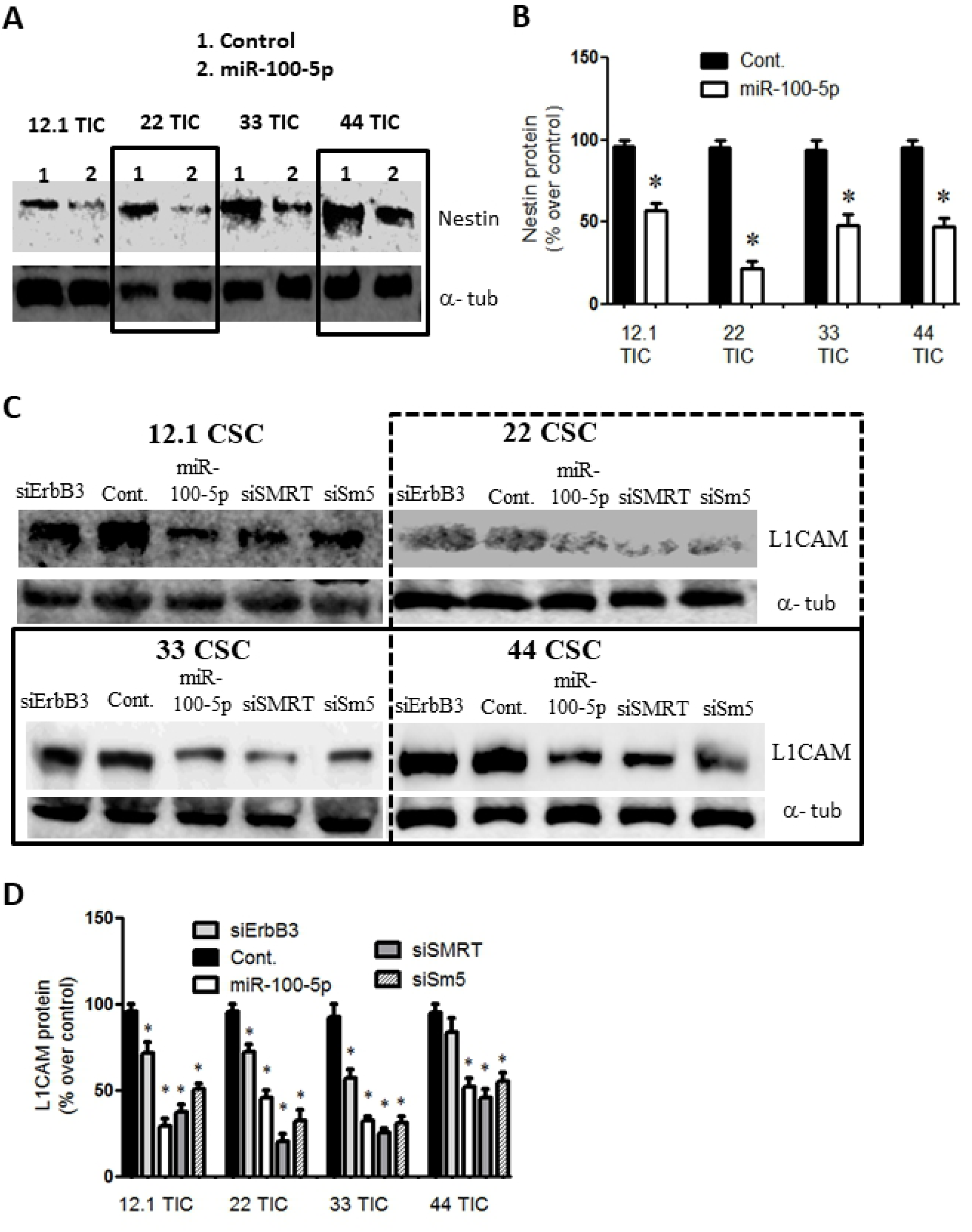
Overexpression of miR-100-5p reduces stem cell markers. (A) Immunoblot showing a reduction in the stem cell marker nestin, following the overexpression of pre-miR-100 or of miR-100-5p but not of the control miRNA. The boxes are included to make the blot easier to interpret. (B) Quantification of protein levels in panel (A) represent reduction of 40–80% as compared to control. (C) Immunoblot showing the reduction in protein levels of the stem cell marker L1CAM when GBM TICs were transfected with pre-miR-100 (miR-100-5p), siSMRT, or Sm5 (siSMARCA5). This reduction was not observed after transfection with the control miRNA. The boxes are included to make the blot easier to interpret. (D) Quantification of the protein levels in panel (C) showing L1CAM reduction of 30–70% compared to the control. Asterisk denotes statistical significance of p<0.05.

### Stable overexpression of pre-miR-100 prevents ErbB2 and ErbB3 expression

GBM tumor cells were transfected with an inducible pre-miR-100-expression vector (resulting in a 3-fold increase in miR-100 levels) and then transplanted into nude mice. Immunohistochemistry was carried out on mouse brain paraffin-embedded slides 16 days after transplantation. The results showed that the expression of ErbB2 and ErbB3 was significantly lower in mice bearing GBM cells transfected with the pre-miR100-expression vector than in mice transplanted with GBM cells transfected with the control vector (Fig. 6 A, B; P<0.05; n = 8).

**Figure 6:**
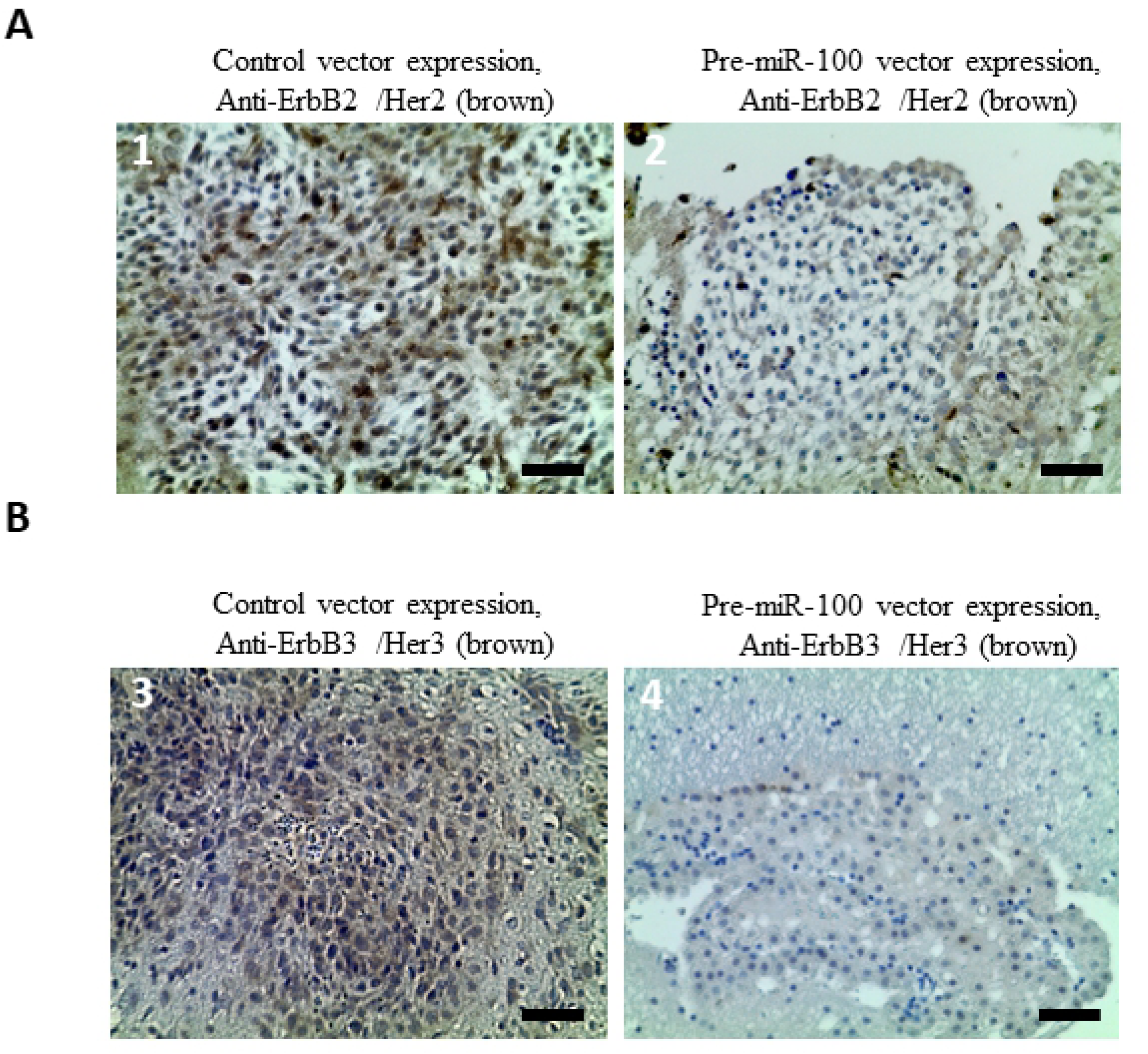
Overexpression of pre-miR-100 decreases the expression of ErbB2 and ErbB3 *in vivo*. (A) Mouse NOD-SCID GBM xenografted brain tissue stably overexpressing pre-miR-100 exhibits less ErbB2 (Her2) expression than the control tissue overexpressing the vector alone. The darker the brown staining, the higher expression of Her2 antigen. The more positive stain is scored with more pluses. The pre-miR-100 group in picture 2, is scored with one plus (+), and the control tissue is scored with 3 pluses (+ + +) in picture 1. (B) GBM xenograft tissue stably overexpressing pre-miR-100 exhibits less ErbB3 (Her3) expression than control tissue overexpressing the vector alone. The pre-miR-100 group is scored with questionable positivity (+/-) in picture 4, and the control group is scored with two pluses (+ +) in picture 3. Scale bar: 50 µm.

## Discussion

We previously reported that the down-regulation of miR-100, and the subsequent reduction in levels of its target SMRT/NCRO2, can contribute to GBM tumorigenicity [13]. We also showed previously that restoring miR-100 levels reduces GBM tumorigenicity and extends the survival of animals bearing orthotopic GBM xenografts [13]. Our results here demonstrate that miR-100 overexpression, by using pre-miR-100 to replenish miR-100 levels in GBM cells, activates the cell cycle inhibitor p21 and inhibits pathways that are important for cell survival and proliferation (e.g., AKT, ERK, and STAT3), and targets SMARCA5 and ErbB3 (Her3).

We observed lower abundance of miR-100-5p in patient-derived GBM TICs compared to human neural stem cells, and found that transfection with pre-miR-100 decreased the proliferation and viability of GBM TICs. This suggests that miR-100 probably plays an important role in limiting GBM growth by modulating one or more of its targets, in addition to SMRT/NCRO2.

By further refining the bioinformatics analysis of miR-100-5p targets, we identified SMARCA5 (SWI/SNF-related matrix-associated actin-dependent regulator of chromatin subfamily A member 5) as a target of miR-100-5p. When we combined the target predictions provided by the microRNA.org, PICTAR, and TARGETSCAN algorithms, only 18 targets were identified in common (Fig. 1C). Of these, SMARCA5 was the only one with a high score and low energy (Table 1). SMARCA5 is reported to be highly expressed in proliferating stem and progenitor cells, and its activity disappears or is reduced considerably in terminally differentiated cells [27, 28]. Thus, we investigated whether SMARCA5 also plays a role in the down-regulation of miR-100-5p in GBM tumors and in the subsequent effect on GBM growth. Luciferase reporter assays confirmed that SMARCA5 is a target of miR-100 (Fig. 1G). In addition, we have demonstrated that miR-100 expression reduces the expression of SNF2H (the *SMARCA5*-encoded protein) in GBM TICs by approximately 50% (Fig. 1G). This finding suggests that miR-100-5p silences the *SMARCA5*-encoded transcript and that the SNF2H protein might play a role in GBM tumorigenesis.

It was reported that SMARCA5 interacts with HDAC3 [29, 30] which modulates AKT. It is also known that AKT controls cell survival and proliferation [31, 32]. In our study, we observed that the knockdown of SMRT or of SMARCA5, or overexpression of pre-miR-100, reduced the phosphorylation of AKT and ERK in GBM TICs. ErbB (Her) family proteins are known to act upstream of both the AKT and ERK pathways [33, 34]. The knockdown of SMRT or SMARCA5, or transfection with pre-miR-100, significantly decreased ErbB3 (Her3) protein levels in GBM TICs. However, the precise mechanism by which ErbB3 protein levels are reduced is not clear because ErbB3 lies upstream of the AKT and ERK pathways. Interestingly, our bioinformatics analysis revealed that ErbB3 is a target of miR-100-3p, the less predominant form of miR-100 that is released from pre-miR-100. In this work, ErbB3 3’UTR luciferase vector experiments confirmed this miR-target relationship (Fig. 3E).

Recent studies have shown that *SMARCA5* needs to be highly expressed for stem cell self-renewal [35] and that HDAC3 is required for the function of the *SMARCA5*-encoded protein SNF2H [30]. Furthermore, SMRT is essential for HDAC3 activity [36-38], and the induction of a DNA-damage response by SMRT inhibition leads to increased p21 levels [39-41]. Consistent with this finding, we observed that p21 protein levels were increased by 100% in GBM TIC lines transfected for miR-100-5p overexpression, or with transfected with an siRNA that silences SMRT relative to control cells (Fig. 4C). This p21 induction resulting from miR-100 activity suggests that GBM TIC proliferation will be inhibited. There is an inverse relationship between p21 levels and STAT3 activation [42-44], and our results showed that the overexpression of either pre-miR-100 or miR-100-5p in GBM TICs almost completely blocked STAT3 phosphorylation (Fig. 4A, B). This suggests that miR100-mediated STAT3 inhibition can exert downstream effects on GBM growth. It worth noting that Tyr705-STAT3 works under interaction of mTOR-STAT3-Notch signaling pathway known to inhibit glioma growth and regulate stem cells [45, 46].

The Notch1 pathway regulates SMRT activity which is known to be involved in cancer stemness [47, 48]. Further, SMARCA5 inhibition stimulates differentiation [49, 50] in GBM TICs which were tested for the loss of stem cell markers. The protein levels of both nestin and L1CAM were significantly lower in GBM TIC cells overexpressing miR-100-5p than in control cells (Fig. 5A and D). As expected, knockdown of SMARCA5 and SMRT also diminished L1CAM protein levels (Fig. 5D). The inhibition of L1CAM (CD171) was previously reported to decrease DNA damage repair (ATM) in GBM TICs through inhibition of NBS1 which activates ATM [51]. This suggests that the loss of miR-100-5p in GBMs plays a role in maintaining the ‘stem cell-like’ status of GBM TICs. Similarly, miR-100 was found to target the SMARCA5-encoded protein SNF2H in adenocarcinoma cells [52]. Thus, SMARCA5 may partially play an important role in cancer by maintaining the de-differentiated state of cancer stem-like cells.

Mice implanted with GBM TICs that were transfected with stable pre-miR-100 expression or inducible pre-miR-100 expression vectors showed tumor xenograft immunostaining that revealed decreased ErbB2 (Her2) and ErbB3 (Her3) protein levels in the pre-miR-100 group compared to the control group (Fig. 6A, B).

This study provides evidence that miR-100 indirectly inhibits three major pathways (STAT3, AKT, and ERK) in GBM TICs. STAT3 is inhibited by the up-regulation of p21 that results from the miR-100-mediated down-regulation of SMRT and/or SMARCA5. AKT and ERK are inhibited by the suppression of either HDAC or ErbB3. The proliferation of GBM TICs was markedly reduced, and differentiation was induced following transfection with pre-miR-100.

In conclusion, our studies show that miR-100 targets many pathways that might contribute to tumorigenicity. A recent randomized clinical trial showed that miR-100 is also responsible for vitamin D-induced tumor suppression in primary prostate cancer [53]. Furthermore, many studies have linked miR-100 with sensitizing tumors to radiotherapy [12, 54, 55]. Findings reported in this study suggest that altering miR-100 levels and its downstream effects (possibly with pre-miR-100 or analogs) might have potential clinical applications in GBM treatment strategies.

## Acknowledgments

We appreciate support from Saudi Ministry of National Guard RC13/258/R (King Abdullah Int’l Medical Research Center, KAIMRC) to BMA, and NIH T32GM007507, UL1RR025011, RC4AA020476, NCI HHSN261201000130C, P30CA014520 grants, the Wisconsin Partnership Program core grant support from Center for Stem Cell and Regenerative Medicine, from the University of Wisconsin (Graduate School and Dept. of Neurological Surgery), and the HEADRUSH Brain Tumor Research Professorship and Roger Loff Memorial Fund for GBM Research to JSK. The funders had no role in study design, data collection and analysis, decision to publish, or preparation of the manuscript.

